# SMAD2 Promotes Myogenin Expression and Terminal Myogenic Differentiation

**DOI:** 10.1101/2020.07.28.225888

**Authors:** Émilie Lamarche, Hamood AlSudais, Rashida Rajgara, Dechen Fu, Saadeddine Omaiche, Nadine Wiper-Bergeron

## Abstract

SMAD2 is a transcription factor whose activity is regulated by members of the Transforming Growth Factor beta (TGFβ) superfamily. While activation of SMAD2 and SMAD3 downstream of TGFβ or myostatin signaling is known to inhibit myogenesis, we find that SMAD2 in the absence of TGFβ signaling promotes terminal myogenic differentiation. We find that during myogenic differentiation, SMAD2 expression is induced. Knockout of SMAD2 expression in primary myoblasts did not affect the efficiency of myogenic differentiation but produced smaller myotubes with reduced expression of the terminal differentiation marker myogenin. Conversely, overexpression of SMAD2 stimulated myogenin expression, and enhanced both differentiation and fusion, and these effects were independent of classical activation by the TGFβ receptor complex. Loss of *Smad2* in muscle satellite cells *in vivo* resulted in decreased muscle fiber caliber and impaired regeneration after acute injury. Taken together, we demonstrate that SMAD2 is an important positive regulator of myogenic differentiation, in part through the regulation of *Myog*.

**SUMMARY STATEMENT:** TGFβ and SMAD2 are normally associated with inhibition of myogenesis. Our work identifies a pro-myogenic role for SMAD2 during terminal myogenic differentiation, through regulation of *Myog* and *Klf4* expression.

## INTRODUCTION

After injury, skeletal muscle can regenerate due to the presence of myogenic precursor cells, called satellite cells, found between the muscle fiber sarcolemma and the basal membrane (Mauro, 1961). Satellite cells are a heterogeneous population characterized by both their histological location and their expression of Paired box protein 7 (PAX7). Normally quiescent, satellite cells are activated following muscle injury to proliferate and differentiate into myocytes that can fuse to one another to form myofibers, or to damaged myofibers to repair them (Charge and Rudnicki, 2004). Adult skeletal myogenesis is a well-organized process governed by the induction and expression of transcription factors known as the myogenic regulatory factors (MRFs). The MRFs, including MYF5, MYOD, myogenin (MYOG) and MRF4, are part of the basic helix-loop-helix family of transcription factors, which bind E-boxes found in many myogenic promoters, and are known for their ability to convert non-myogenic cells to the myogenic lineage by upregulating muscle specific genes (Wang and Rudnicki, 2012). MYF5 and MYOD are important myogenic commitment factors while myogenin and MRF4 are induced later in differentiation and are necessary for the development of mature muscle. MYOD^-/-^ animals have normal skeletal muscle (Rudnicki et al., 1992), a loss compensated by prolonged MYF5 expression (Megeney et al., 1996). MYF5^-/-^ animals die shortly after birth of respiratory failure due to a malformed rib cage, but similar to MYOD^-/-^ animals, they have relatively normal skeletal muscle with unchanged expression of MYOD, myogenin and MRF4 (Braun et al., 1992). Myogenin^-/-^ mice die perinatally from respiratory failure due to a severe reduction in all skeletal muscle, characterized by an abundance of mononucleated cells and rare myofibers, with a failure to induce the contractile protein myosin heavy chain (MyHC), suggesting these myoblasts are committed to the myogenic lineage but fail to differentiate and form myofibers (Hasty et al., 1993; Nabeshima et al., 1993). MRF4^-/-^ mice have normal expression of muscle-specific genes, suggesting that myogenin has the crucial role in myogenic differentiation (Hasty et al., 1993).

Many signaling pathways regulate myogenic differentiation, including members of the Transforming Growth Factor beta (TGFβ) superfamily, of which TGFβ is a potent inhibitor (Liu et al., 2001; Liu et al., 2004). Both TGFβ and family member myostatin activate the transcription factors SMAD2 and SMAD3 through receptor-dependent phosphorylation of their C-termini, promoting their interaction with SMAD4, nuclear localization and interaction with response elements in the promoters and enhancers of target genes (Shi and Massague, 2003). As such, SMAD2 and SMAD3 are largely considered to mediate an anti-myogenic arm of TGFβ family signaling. However, despite the inhibitory role of myostatin and TGFβ signaling on myogenesis, loss of SMAD3 expression in myoblasts impairs myogenic differentiation (Ge et al., 2011; Ge et al., 2012), suggesting that SMAD3 has some pro-myogenic functions. In accordance with this, retinoic acid treatment, which promotes myogenic differentiation, can stimulate SMAD3 expression and rescue myogenesis in the presence of TGFβ, at least in part through a physical interaction between SMAD3 and C/EBPβ, a transcriptional factor that is present in undifferentiated cells and acts to inhibit myogenic differentiation (Lamarche et al., 2015).

SMAD2 and SMAD3 are highly conserved, are activated similarly and bind the same DNA response element in target promoters, however there is increasing evidence that these two transcription factors have divergent roles *in vivo* with functions beyond classical TGFβ signaling. While SMAD3-null mice are viable and fertile (Yang et al., 1999; Zhu et al., 1998), deletion of SMAD2 is embryonic lethal as embryos fail to gastrulate and induce mesoderm (Nomura and Li, 1998; Waldrip et al., 1998; Weinstein et al., 1998), suggesting that during development, SMAD2 is of greater importance. While the role of SMAD3 in myogenesis has been investigated, little is known about the specific contribution of SMAD2 to myogenic differentiation. Herein, we developed a conditional SMAD2 knockout mouse where SMAD2 expression is abolished in PAX7^+^ muscle myogenic precursor (satellite) cells (*Smad2*^SC-/-^). We report that SMAD2 is required for efficient myogenic differentiation in a TGFβ-independent mechanism. In its absence, postnatal muscle growth and muscle regeneration after injury are impaired. We find that SMAD2 regulates the expression of key regulators of myogenesis to promote differentiation.

## RESULTS

### SMAD2 expression promotes myogenic differentiation

To characterize the role of SMAD2 during adult myogenesis, we first quantified *Smad2* mRNA and protein expression in the myogenic cell line C2C12, cultured under growth conditions and after induction to differentiate. *Smad2* mRNA expression was relatively stable during myoblast differentiation (Fig. 1A). Interestingly, SMAD2 protein expression was lower in proliferating myoblasts seeded at low density (low confluency, LC) but increased as culture density increased (high density, HC) and in early myogenic differentiation, coincident with the upregulation of myogenin expression (Fig. 1B,C). After 2 days of differentiation, SMAD2 protein levels returned to levels observed in sub-confluent cells.

**Figure 1.**
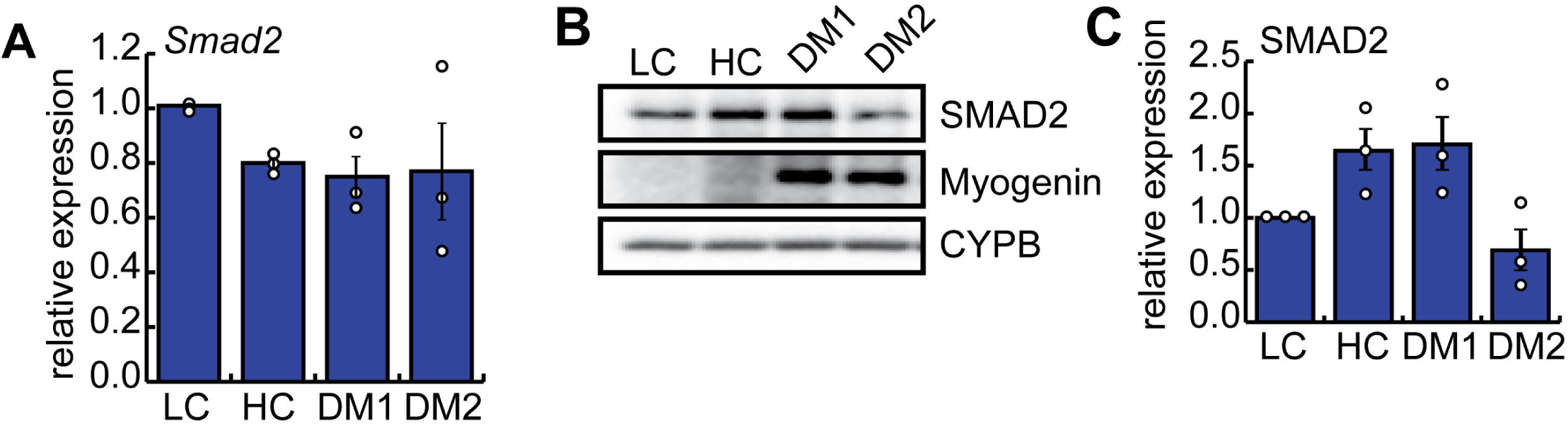
SMAD2 expression is regulated during myogenic differentiation. **(A)** RTqPCR analysis of *Smad2* expression in primary myoblasts isolated from C57BL/6 mice and cultured in growth medium at low confluency (LC), high confluency (HC) or following induction to differentiate in low serum conditions (differentiation medium, DM) for one day (DM1) or 2 days (DM2). Bars are means +STDEV, n=4. **(B)** Representative western blot of SMAD2 and MYOG expression in primary myoblasts under growth conditions and after induction to differentiate for the indicated days. CyPB is a loading control. **(C)** Quantification of SMAD2 and myogenin protein expression from (B) as compared to LC. n=3.

To determine the role of SMAD2 during myogenic differentiation, C2C12 myoblasts were retrovirally transduced to express full length SMAD2 or with empty virus (pLPCX) and overexpression of SMAD2 was confirmed by Western blot and RT-qPCR in proliferating cells (Fig. 2A,B). After differentiation for up to 4 days under low serum conditions, overexpression of SMAD2 increased both the percentage of cells that differentiated as well as the average myotube size (fusion index), suggesting that SMAD2 positively regulates myogenic differentiation. (Fig 2C-E). Analysis of myogenic marker expression revealed that SMAD2 overexpression did not impact *Smad3* expression, but stimulated both *Myog* and *Tmem8c* (myomaker) expression after one day of differentiation (DM 1) (Fig. 2F). Myogenin protein levels were also increased on day 1 of differentiation as compared to control cells (Fig. 2G,H), consistent with enhanced myogenic differentiation and fusion.

**Figure 2.**
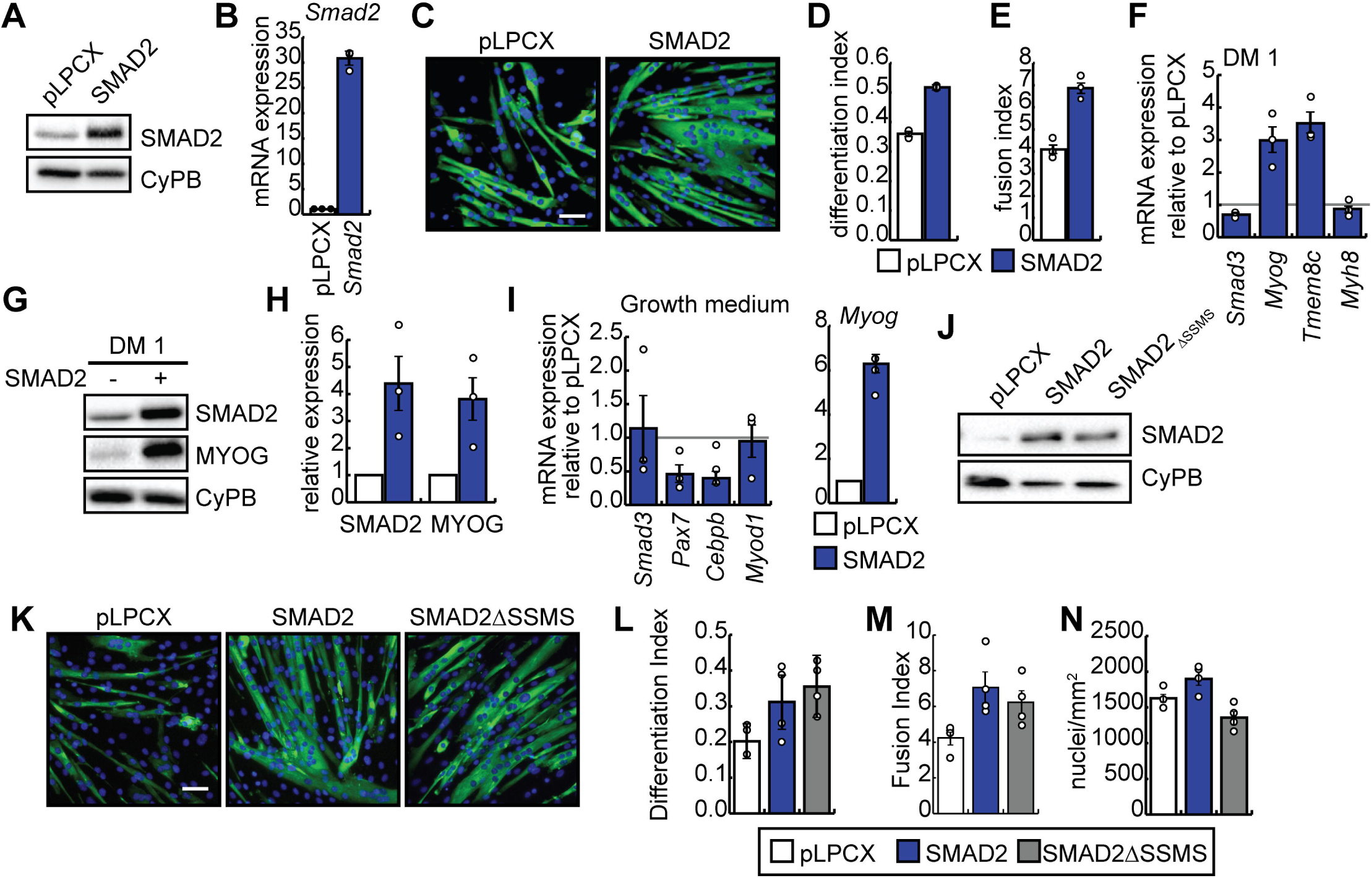
Overexpression of SMAD2 enhances terminal differentiation and myoblast fusion. **(A)** Western blot of SMAD2 expression in C2C12 myoblasts retrovirally transduced to express SMAD2 or with empty virus (pLPCX) and differentiated for 24 hours. Cyclophilin B (CyPB) is a loading control. **(B)** RTqPCR analysis of *Smad2* expression in cells from (A). Bars are means +STDEV. n=3. **(C)** Immunostaining for MyHC (green) in cells transduced as in (A) and induced to differentiate for 4 days. DAPI (blue) counterstains the nuclei. Scale bar = 50 µm. **(D)** Differentiation index (# nuclei in MyHC+ cells/ total nuclei) from cells differentiated as in (C). n=3. **(E)** Fusion index (# nuclei found in MyHC+ cells with 2 or more nuclei/ # myotubes) from cells differentiated as in (C). n=3. **(F)** *Smad3* and myogenic marker mRNA expression in myoblasts transduced as in (A) after induction to differentiate for one day (DM1). n=3. **(G)** Representative western blot of SMAD2 and MYOG expression in myoblasts transduced as in (A) after induction to differentiate for one day (DM 1). Cyclophilin B (CyPB) is a loading control. **(H)** Quantification of western blots represented in (G). n=3. **(I)** RTqPCR analysis of *Smad3, Pax7, Cebpb, Myod1 and Myog* expression in myoblasts transduced as in (A) and cultured in growth medium. Data for *Smad2*-overexpressing cultures is shown as the means relative to controls indicated by the red line. n=3. **(J)** Western blot of C2C12 cells transduced to express SMAD2 or a truncated SMAD2 lacking the C-terminal SSMS motif (SMAD2ΔSSMS). **(K)** Immunostaining for MyHC (green) cells from (J) differentiated for 4 days. Scale bar = 50 µm. **(L)** Differentiation index (# nuclei in MyHC+ cells/ total nuclei) from cells differentiated as in (K). n=3. **(M)** Fusion index (# nuclei found in MyHC+ cells with 2 or more nuclei/ # myotubes) from cells differentiated as in (K). n=3. **(N)** Cell culture density expressed as nuclei/mm^2^ in images used to calculate (L,M). n=3.

Given the positive effect of SMAD2 overexpression on myogenic differentiation, we assessed the expression of markers associated with undifferentiated and differentiated cells under growth conditions (Fig. 2I). As in differentiating cells, *Smad3* levels were not affected by *Smad2* overexpression under growth conditions. However, the expression of *Pax7* and *Cebpb*, two markers associated with the undifferentiated state, was downregulated in cells overexpressing *Smad2* while *Myod1* expression was unaffected (Fig. 2I). Further, *Myog* expression was greatly increased in cells overexpressing *Smad2*, suggesting that SMAD2 overexpression promotes precocious differentiation of myoblasts under growth conditions (Fig. 2I).

Since myoblasts can produce TGFβ ligands and do express TGF receptors, we generated pooled stable C2C12 cell lines expressing full length SMAD2 or a truncated SMAD2 in which the C-terminal SSMS motif, targeted by the activated TGFβ receptor complex, was deleted (SMAD2ΔSSMS) (Fig. 2J). The SMAD2ΔSSMS mutant is not responsive to TGFβ signaling (Choy et al., 2000). Upon differentiation, the SMAD2ΔSSMS was able to enhance myogenic differentiation and fusion similarly to full length SMAD2 (Fig. 2K-M) without impacting cell numbers (Fig. 2N), suggesting that the stimulation of myoblast fusion by SMAD2 does not depend on the presence of the C-terminal SSMS motif and therefore classical TGFβ signaling pathways.

We next characterized the myogenic potential of primary myoblasts deficient for *Smad2*. Primary myoblasts were isolated from the *Smad2*^fl/fl^ mouse (Ju et al., 2006) and retrovirally transduced to express a tamoxifen-regulated Cre recombinase (CreER) (Nishijo et al., 2009). Excision of *Smad2* was achieved by tamoxifen (4OH-TAM) treatment of pooled stable cell lines to generate a *Smad2*-deficient cell line (+TAM), or with vehicle to generate controls (Veh). Tamoxifen treatment resulted in near complete loss of *Smad2* mRNA expression (Fig. 3A). To determine the impact of *Smad2* depletion on myogenic differentiation, TAM- and vehicle-treated cultures were induced to differentiate for 2 days under low serum conditions. While the loss of *Smad2* expression did not negatively impact myogenic differentiation as measured by the percentage of nuclei found within myosin heavy chain-positive (MyHC+) cells (Differentiation index, Fig. 3B,C), myotube maturation was impaired as evidenced by much smaller myotubes in cultures lacking *Smad2* and a reduced fusion index (average # nuclei/myotube) (Fig. 3B,D). The culture density was unaffected by loss of *Smad2* expression (Fig. 3E). While the expression of *Tmem8c* (myomaker) and *Myod1* was highly variable and largely unchanged in tested cells, *Myog* and neonatal myosin heavy chain (*Myh8*), markers of later differentiation, were significantly reduced in *Smad2*-deficient cultures, consistent with impaired terminal differentiation (Fig. 3F). Expression of *Smad3* was unchanged with knockdown of *Smad2*, suggesting that this factor does not increase its expression to compensate for the loss of *Smad2* in myoblasts (Fig. 3F). Western blot analysis revealed that cells lacking SMAD2 also express less MYOG protein as compared to controls (Fig. 3G). Taken together, our data indicates that SMAD2 expression is required for efficient myogenic differentiation, at least in part, through the regulation of myogenin expression.

**Figure 3.**
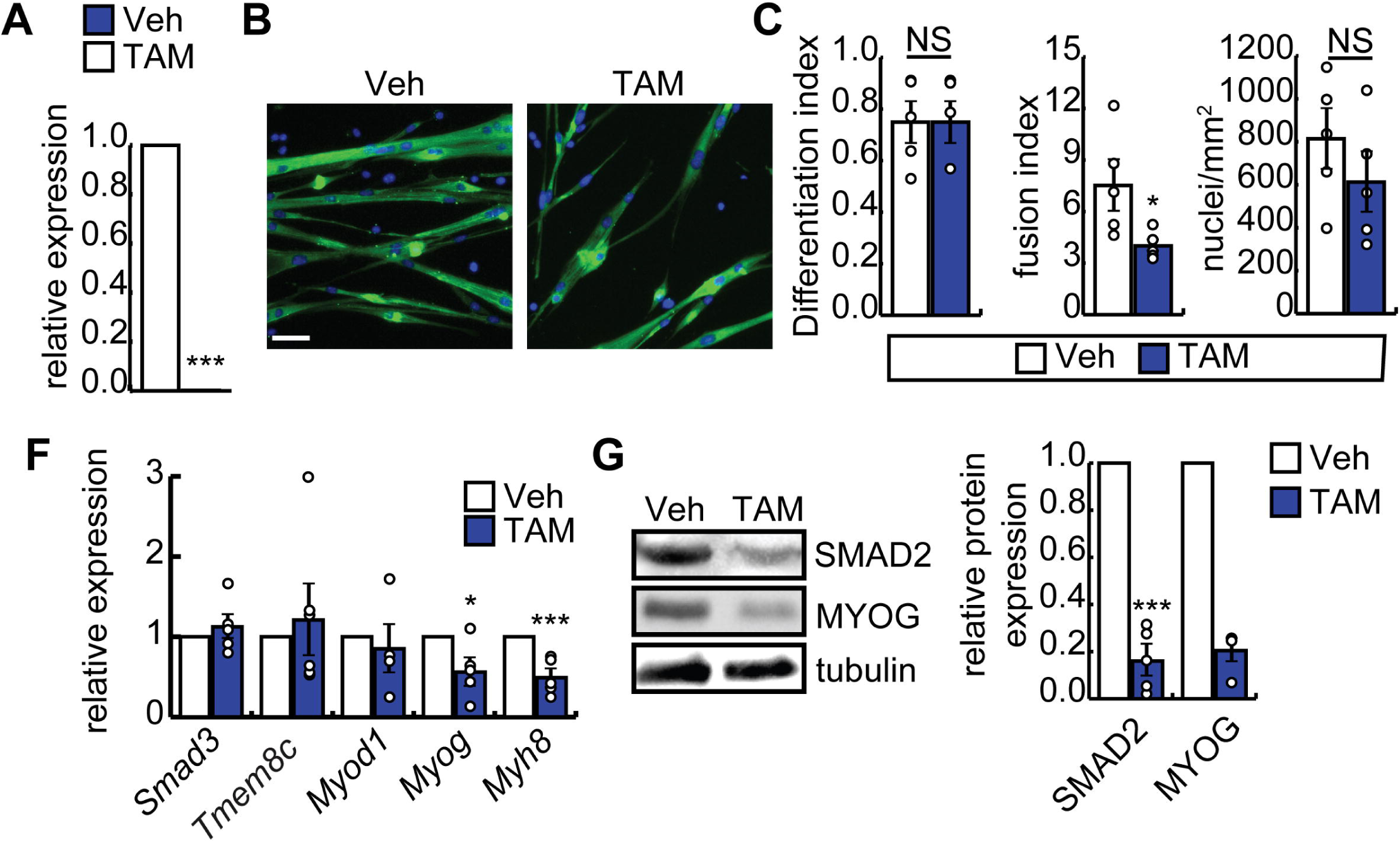
SMAD2 regulates terminal myogenic differentiation. Primary myoblasts isolated from *Smad2*^fl/fl^ mice were retrovirally transduced to express the tamoxifen-inducible Cre recombinase (CreER) and treated with 4-OH tamoxifen (TAM) for 3 days to induce excision or with vehicle (Veh) to generate controls. **(A)** *Smad2* mRNA expression following induction of *Smad2* excision with 4-OH tamoxifen after 48 hours in DM. ***p<0.001, n=5. **(B)** Immunostaining for MyHC expression in TAM-treated and vehicle-treated myoblasts differentiated for 2 days and counterstained with DAPI (blue). Scale bar = 50 µm. **(C)** Differentiation index (# nuclei in MyHC+ cells/ total nuclei) for cells in (B), n=5. NS is not significant. **(D)** Fusion index (# nuclei found in MyHC+ cells with 2 or more nuclei/ # myotubes) from cells differentiated as in (B). *p<0.05, n=5. **(E)** Number of nuclei per mm^2^ counted in (C). n=5. **(F)** RTqPCR analysis of *Smad3, Pax7* and myogenic marker expression in cells transduced and differentiated as in (B). ***p<0.001, n=5. **(G)** SMAD2 and MYOG expression in control and SMAD2-deficient primary myoblasts differentiated as in (B) (left) and quantification of protein expression (right), ***p<0.001, n=5 (SMAD2), n=3 (MYOG).

### Knockdown of SMAD2 inhibits muscle regeneration after acute injury

To determine the *in vivo* role for SMAD2 in the regulation of myogenesis, we generated a conditional knockout model in which *Smad2* is excised in muscle satellite cells by breeding the *Smad2*^fl/fl^ mouse (Ju et al., 2006) with a *Pax7*^CreER/+^ driver line (Nishijo et al., 2009). To assess the myogenic potential of *Smad2*-deficient satellite cells *in vivo*, we treated *Smad2*^fl/fl^*Pax7*^CreER/+^ (*Smad*^SC-/-^) and non-Cre expressing littermates (*Smad2*^fl/fl^*Pax7*^+/+^; WT) with tamoxifen at 6 weeks of age to induce excision of *Smad2*. One week after tamoxifen treatment, we injured the TA muscle in *Smad2*^SC-/-^ and controls with cardiotoxin and assessed the extent of repair seven days after injury (Fig. 4). SMAD2 protein expression in freshly isolated myoblasts from *Smad2*^SC-/-^ and controls confirmed efficient knockdown in this model (Fig. 4A). One week after injury, WT muscle repaired efficiently, regaining a fiber cross-sectional area approximately 50% of uninjured controls (Fig. 4B,C, white bars). *Smad2*^SC-/-^ muscle, however, had impaired regeneration, with fiber cross-sectional areas significantly smaller than those of injured WT mice and uninjured controls (Fig. 4B,C, blue bars). Since regeneration is dependent on PAX7^+^ cells, immunostaining for PAX7 was performed in repairing muscle and uninjured control muscle to assess the population size. The number of PAX7^+^ cells per area was not different between genotypes in the uninjured TA muscles and both underwent expansion following injury (Fig. 4D). However, we did observe a small but significant decrease in the number of PAX7+ cells in *Smad2*^SC-/-^ cardiotoxin-injured muscle. To determine if regeneration was impaired or simply delayed, we repeated the injury experiment and harvested TA muscle 14 days post-injury. The remaining hindlimb muscles were digested for satellite cell isolation to confirm inactivation of SMAD2 (Fig. 4E). At this time point, the regenerating myofiber cross-sectional area was indistinguishable from WT TA muscle, suggesting that loss of SMAD2 causes a delay in muscle regeneration (Fig. 4F,G). The number of PAX7^+^ cells was unchanged from control in both the injured and uninjured muscle, suggesting the mild reduction in satellite cell numbers is unlikely to underlie impaired regeneration (Fig. 4H). Interestingly, the number of Myogenin-positive nuclei was increased in *Smad2*^SC-/-^ muscle, suggesting that regeneration is delayed in the absence of *Smad2* (Fig. 4I).

**Figure 4.**
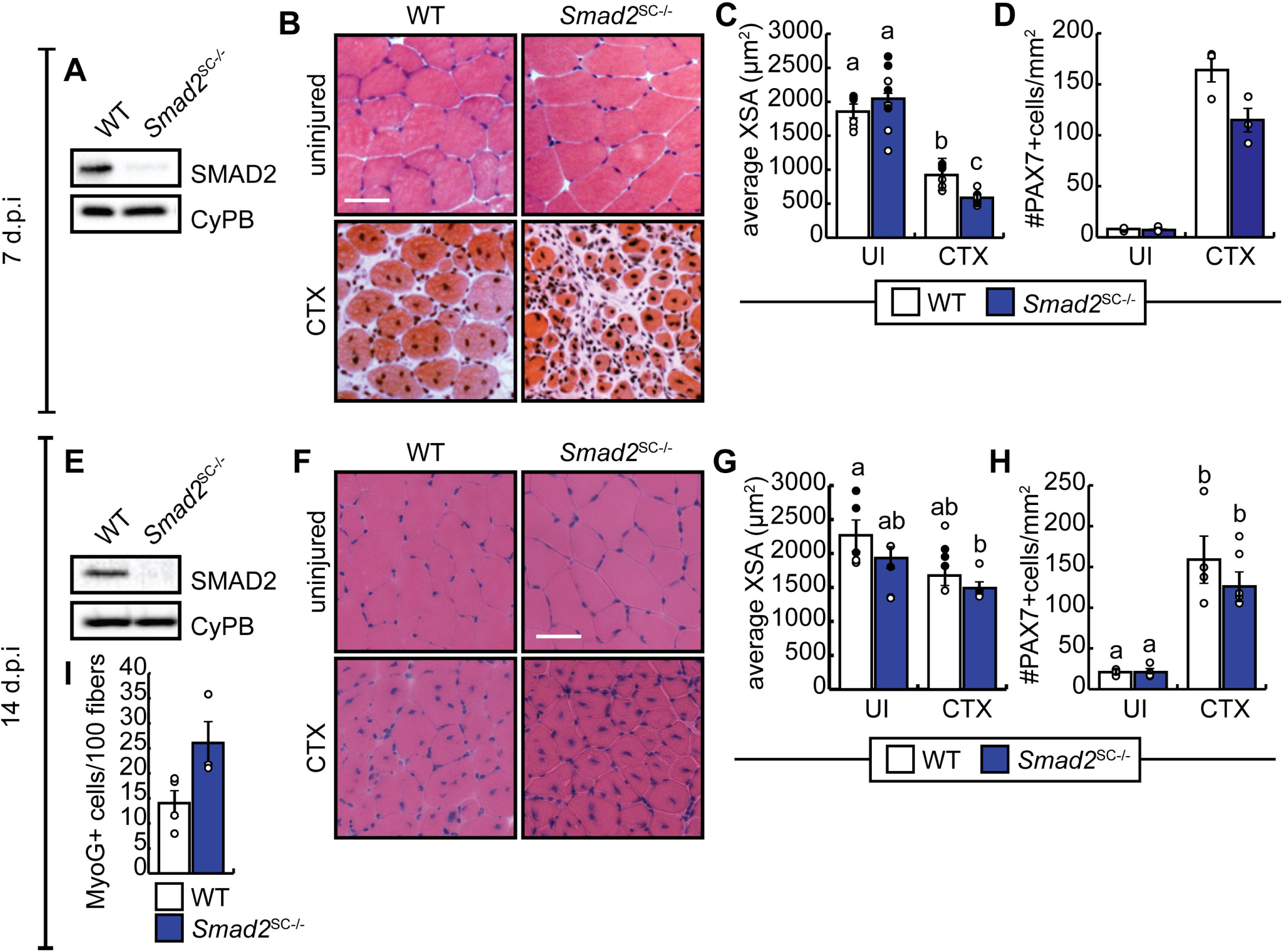
SMAD2 is required for efficient muscle regeneration. **(A)** Representative western blot of SMAD2 protein expression in isolated satellite cells from WT and *Smad2*^SC-/-^ hindlimb 7 days after injury with cardiotoxin to the left TA muscle (7 d.p.i.). Cyclophilin B is a loading control. **(B)** Representative images of cardiotoxin (CTX)-injured and uninjured TA muscle sections from WT and *Smad2*^SC-/-^ mice after repair for 7 days. Scale bar = 50 µm. **(C)** Average cross-sectional area of muscle fibers in WT and *Smad2*^SC-/-^ mice injured as in (B). n= 8 pairs. Black dot data points are male mice and white dots represent female mice. **(D)** Number of PAX7+ cells per area of uninjured and injured TA from (B), n=3. **(E)** Representative western blot of SMAD2 protein expression in isolated satellite cells from WT and *Smad2*^SC-/-^ hindlimb 14 days post-injury with cardiotoxin to the left TA muscle (14 d.p.i). Cyclophilin B is a loading control. **(F)** Representative images of cardiotoxin (CTX)-injured and uninjured TA muscle sections from WT and *Smad2*^SC-/-^ mice after repair for 14 days. Scale bar = 50 µm. **(G)** Average cross-sectional area of muscle fibers in WT and *Smad2*^SC-/-^ mice injured as in (F). Means indicated with different letters are significantly different from one another at a minimum cut-off of p<0.05, n= 5 pairs. Black dot data points are male mice and white dots represent female mice. **(H)** Number of PAX7+ cells per area of uninjured and injured TA from (F), n=5. **(I)** Number of MYOG+ cells per area of injured TA from (F). n=5.

### Loss of SMAD2 in utero perturbs post-natal fiber growth

Knockout of *Smad2* is embryonic lethal, with perturbed formation of mesoderm, and thereby skeletal muscle (Nomura and Li, 1998; Waldrip et al., 1998; Weinstein et al., 1998). To assess the contribution of SMAD2 to the extensive satellite cell differentiation occurring during the post-natal period, *Smad2*^fl/fl^*Pax7*^wt/wt^ females were bred to *Smad2*^fl/fl^*Pax7*^CreER/wt^ males to generate *Smad2*^fl/fl^*Pax7*^CreER/+^ (*Smad2*^SC-/-^) and *Smad2*^fl/fl^*Pax7*^+/+^ (WT) progeny mice and *Smad2* was excised *in utero* by gavage of the pregnant dams at embryonic day 15.5 (E15.5) with tamoxifen. Pups were subsequently sacrificed at post-natal day 21 (P21) and tibialis anterior muscles were dissected and flash frozen for histological analysis and PAX7 immunostaining. The remaining hindlimb muscles were digested for satellite cell isolation to confirm loss of SMAD2 expression (Fig. 5A). Hematoxylin and Eosin staining and analysis of the cross-sectional area (XSA) revealed significantly smaller myofibers at P21 in *Smad2*^SC-/-^ muscle as compared to WT (Fig. 5B,C). Further, the TA muscle of *Smad2*^SC-/-^ had an average of approximately 3000 fibers while WT TA had approximately 2500 (Fig. 5D). Immunostaining revealed no difference in the percentage of PAX7+ cells in the TA muscle at P21 of *Smad2*^SC-/-^ mice as compared to WT muscle sections (Fig. 5E), suggesting that the smaller fiber calibre was not due to reduced satellite cell numbers.

**Figure 5.**
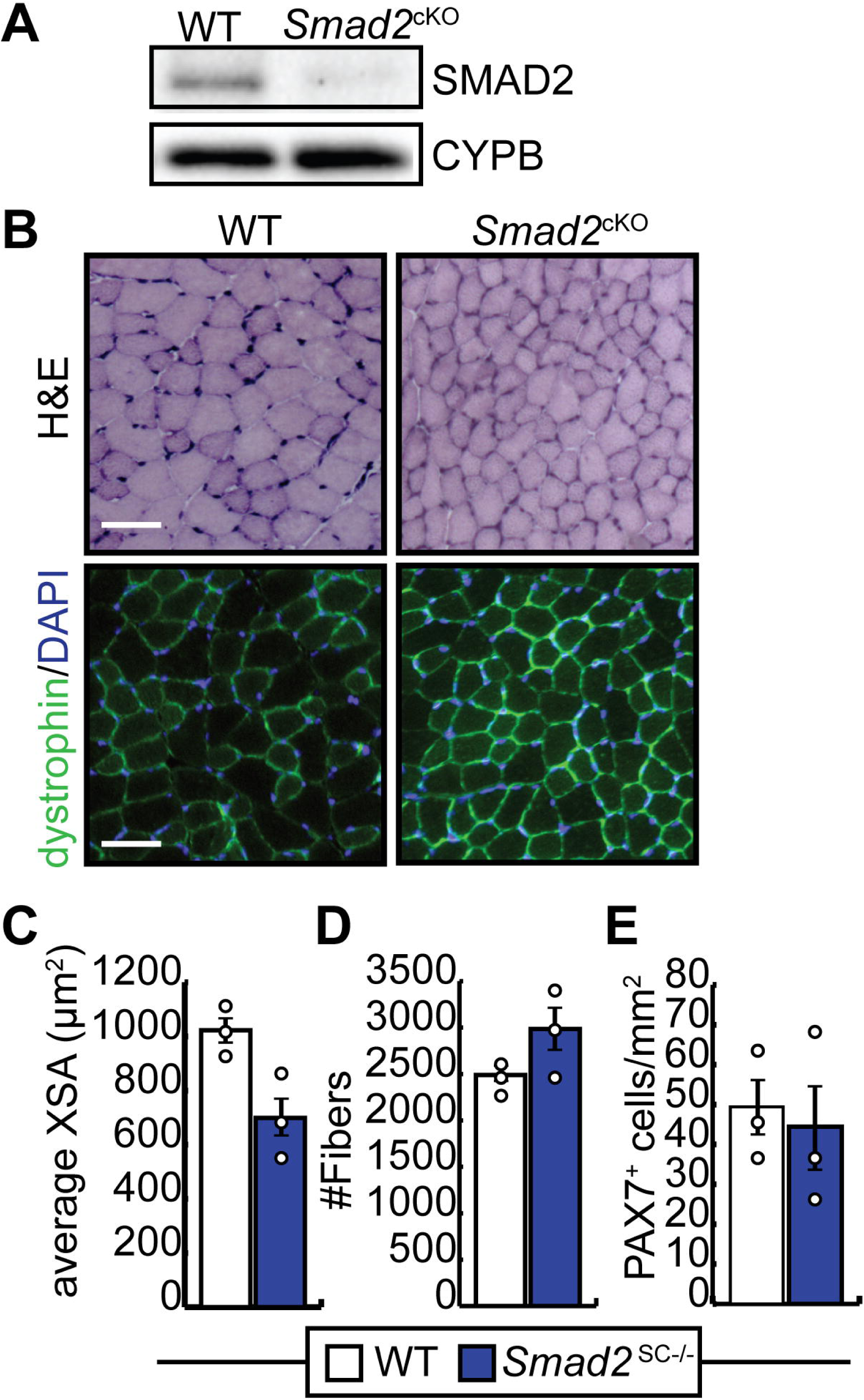
SMAD2 regulates fiber size after birth. **(A)** Western analysis of SMAD2 and CYPB (loading control) from satellite cells isolated from *WT* and *Smad2*^SC-/-^ pups on postnatal day 21 (P21) produced from *Smad2*^fl/fl^*Pax7*^wt/wt^ pregnant dams bred with *Smad2*^fl/fl^*Pax7*^CreER/wt^ males and gavaged with tamoxifen at E15.5. **(B)** Hematoxylin and eosin (H&E) staining (top) and anti-dystrophin immunostaining (bottom) of muscle sections from WT and *Smad2*^cSC-/-^ mice at P21; n=3 pairs. Scale bar = 50 µm **(C)** Average cross-sectional areas (XSA) calculated from muscle sections as in (A); n=3 pairs. **(D)** Average number of fibers in the TA muscle of WT and *Smad2*^cKO^ pups from muscle sections as in (A). **(E)** Number of PAX7^+^ cells per area in muscle sections from (B). Graphs are means + STDEV.

### SMAD2 promotes myoblast fusion

C2C12 myoblasts overexpressing SMAD2 have enhanced fusion and fewer unfused MyHC+ myocytes in culture after differentiation (Fig. 2F, 6A). Since fusion is regulated by numerous factors and myomaker expression was increased by SMAD2 overexpression (Fig. 2F), we examined the expression of the known pro-fusogenic gene *Klf4* which promotes fusion through upregulation of *Npnt* (Sunadome et al., 2011). After 24 hours of differentiation (DM 1), KLF4 protein expression was increased in cells overexpressing SMAD2 (Fig. 6B). To confirm that SMAD2 could directly regulate transcription from the *Klf4* promoter, we performed a reporter assay using a (−1481/+45) *Klf4*-luciferase construct in C2C12 myoblasts. Expression of SMAD2 in C2C12 cells increased *Klf4* promoter activity by approximately 5-fold, suggesting that SMAD2 can directly regulate transcription of *Klf4* in myoblasts (Fig. 6C). We next explored the *Klf4* regulatory region, and identified using published SMAD3 ChIP-seq data (Mullen et al., 2011) coupled to motif analysis, putative SMAD binding elements in the *Klf4* promoter (pro) and enhancer regions (−10kb). Chromatin immunoprecipitation revealed that SMAD2 occupies the *Klf4* regulatory region under differentiation conditions, and thus likely directly regulates *Klf4* expression (Fig. 6D). Consistent with this, expression of the pro-fusogenic gene nephronectin (*Npnt*), a known KLF4 target gene, was enhanced by SMAD2 overexpression (Fig. 6E).

**Figure 6.**
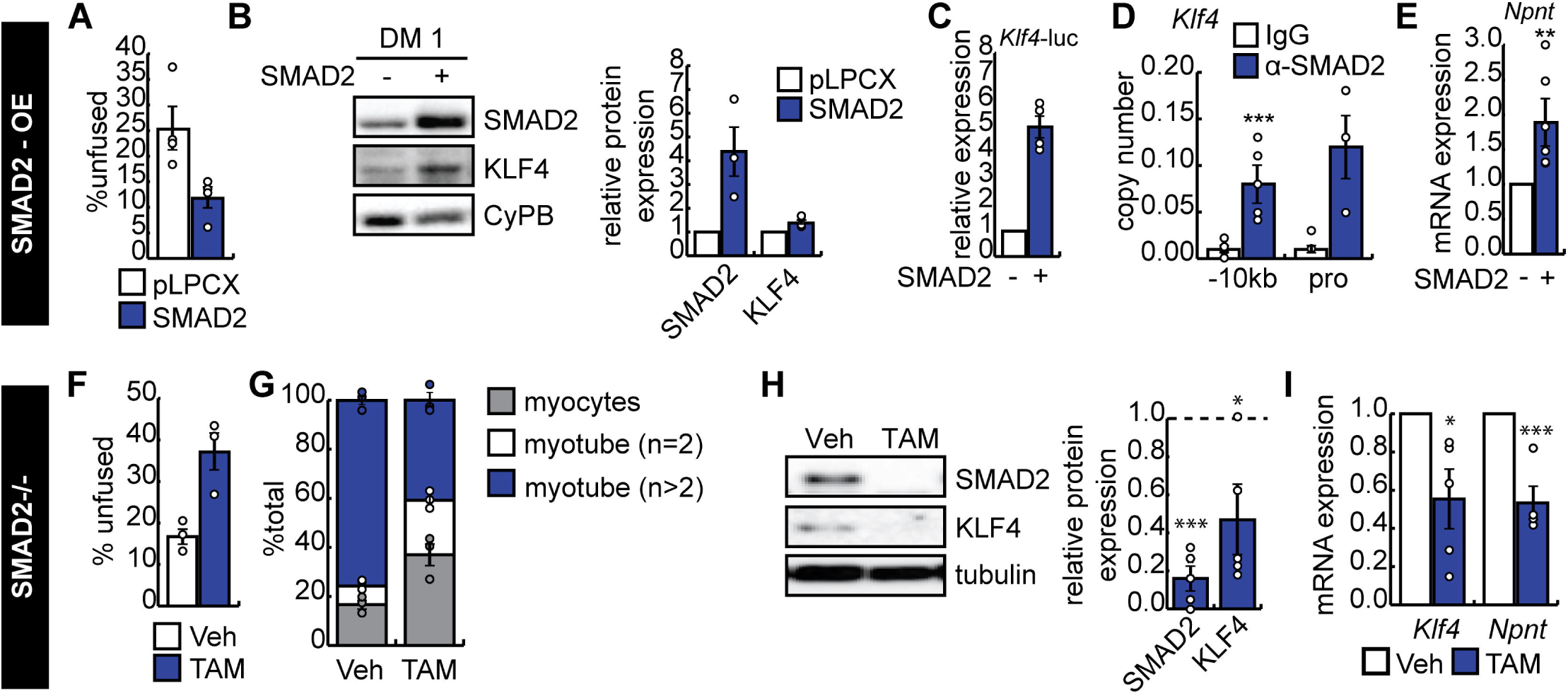
SMAD2 regulates KLF4 expression to promote myoblast fusion. **(A)** Percent of mononucleated MyHC+ cells in cultures of C2C12 cells transduced to express SMAD2 or with empty virus (pLPCX) and induced to differentiate for 4 days. n=4. **(B)** Representative western blot and protein expression quantification of KLF4 expression in differentiating SMAD2-overexpressing primary myoblasts differentiated for one day (DM 1). n=3. **(C)** Luciferase reporter assay to measure *Klf4* promoter activity in the presence or absence of ectopic SMAD2, shown relative to reporter alone and corrected for transfection efficiency. n=4. **(D)** Chromatin immunoprecipitation of SMAD2 recruitment to the *Klf4* promoter and -10 kb upstream region containing putative SMAD2 motifs in primary myoblasts isolated from C57BL/6 mice and differentiated for 1 day. Data is shown as copy numbers (white bars) as compared to pulldown with type-matched IgG as a control (black bars). ***p<0.001, n=6 (−10kb) and n=3 (pro). **(E)** *Npnt* expression in cells cultured as in (B). n=5. **(F)** Percent of mononucleated MyHC+ in control (veh) and *Smad2*^-/-^ primary myoblasts (TAM) after differentiation for 48 hours. n=3. **(G)** Percent of myocytes, nascent myotubes (myotubes with 2 nuclei) and myotubes (>2 nuclei) in cultures from (F). *p<0.05, **p<0.01, ***p<0.001, n=5. **(H)** Representative western blot and quantification relative to controls (dotted line) of KLF4 expression from cultures in (F). *p<0.05, ***p<0.001, n=5. **(I)** *Klf4* and *Npnt* mRNA expression in primary myoblasts differentiated as in (F), *p<0.05, ***p<0.001, n=5.

In *Smad2*-deficient primary myoblasts, fusion was reduced, and we observed an increase in the percentage of unfused MyHC+ cells (Fig. 6F). Further, we found that the number of nascent myotubes, with a total of two nuclei, was also increased as compared to controls, with a concomitant reduction in the percentage of larger myotubes (Fig. 6G). After 12 hours of differentiation, *Smad2*-deficient cells had significantly reduced KLF4 protein expression and *Klf4* mRNA expression (Fig. 6H,I) consistent with the observed phenotype. Further, we observed reduced expression of the KLF4 target gene *Npnt* (Fig. 6I).

Given that *Klf4* is a transcriptional target of SMAD2 in myoblasts and that fusion is enhanced in cells overexpressing SMAD2 and perturbed in cells lacking SMAD2, we hypothesized that SMAD2 acts through KLF4 to enhance fusion. To test this, we retrovirally transduced WT or *Smad2*-deficient primary myoblasts to express KLF4 or with empty virus and induced their differentiation (Fig. 7). Introduction of KLF4 did not affect the differentiation of myoblasts of either genotypes, but enhanced the fusion of myoblasts isolated from WT cells (Fig. 7A-C). However, KLF4 overexpression in *Smad2*-deficient cells failed to rescue fusion (Fig. 7A,C). Consistent with these findings, while *Klf4* mRNA expression was increased in *Smad2*-deficient cells overexpressing *Klf4*, the expression of the downstream target *Npnt* was not rescued in the absence of SMAD2 6 hours after induction to differentiate in low-serum conditions (Fig. 7D). Similarly, *Myog* expression, which was reduced at this time point in *Smad2*-deficient cells, was not rescued by addition of KLF4 (Fig. 7D). Since myogenin expression was influenced by SMAD2 in our experiments, we verified if SMAD2 could regulate *Myog* expression directly using a *Myog*-luc construct in a reporter assay in the presence of ectopic SMAD2 (Fig. 7E). While the addition of SMAD2 did not increase reporter activity, KLF4 was found to be a potentiator of *Myog* promoter activity (Fig. 7E). To determine if SMAD2 was required for KLF4 action on the *Myog* promoter, C2C12 myoblasts were transduced to express a shRNA construct directed against *Smad2* (shSmad2) and the reporter assay was repeated in the presence of KLF4. Addition of SMAD2 enhanced transcription by KLF4 from the *Myog* promoter in the shSmad2 cells suggesting that SMAD2 and KLF4 cooperate to regulate *Myog* expression (Fig. 7F). Next, KLF4 and SMAD2 recruitment to the *Myog* promoter was assessed by chromatin immunoprecipitation, and both were found to be enriched (Fig. 7G). As an interaction between KLF4 and SMAD2 was reported in vascular smooth muscle cells (Li et al., 2010), we immunoprecipitated SMAD2 from C2C12 cells, which efficiently coprecipitated KLF4, suggesting that these proteins interact in myoblasts (Fig. 7H).

**Figure 7.**
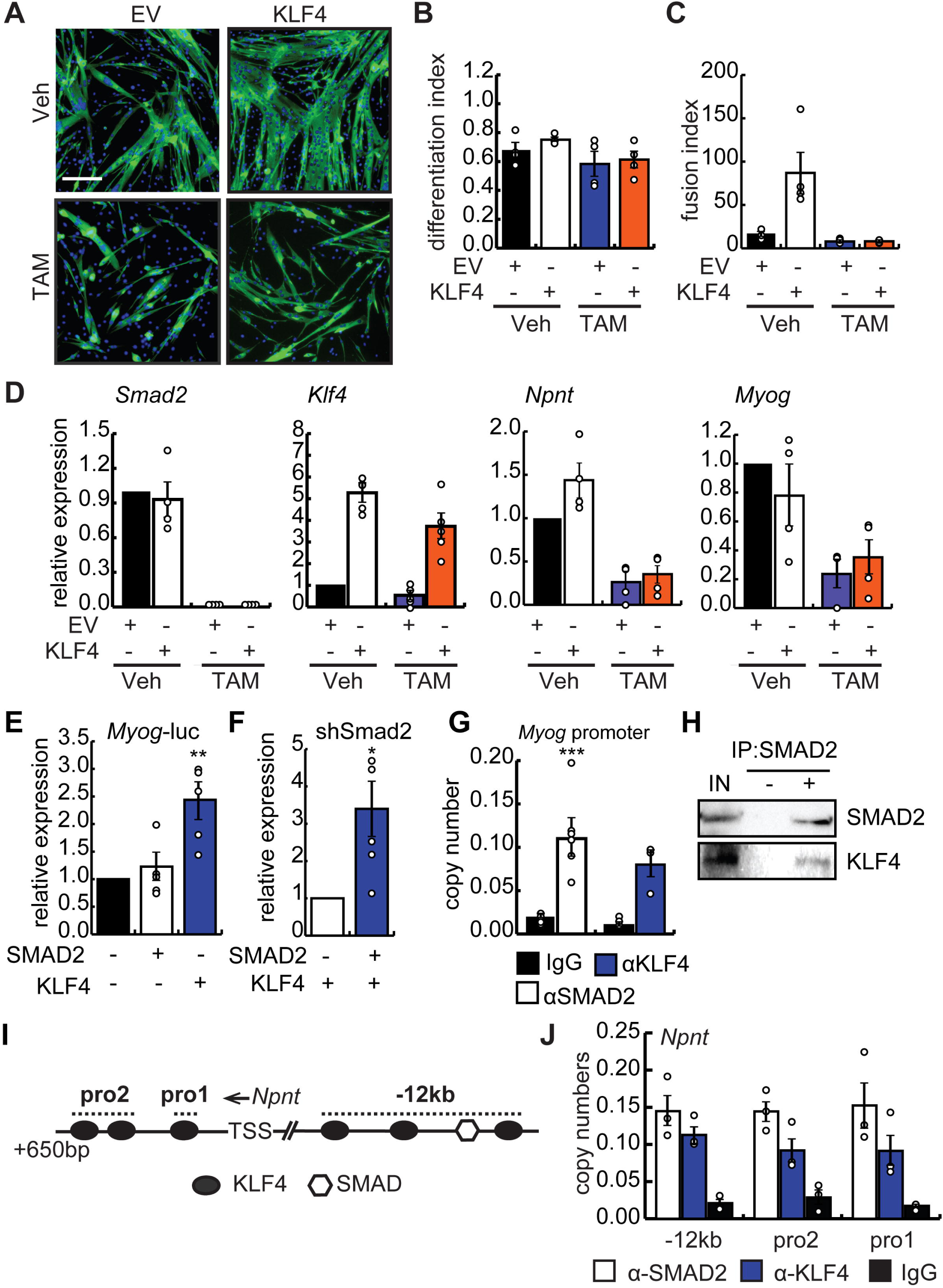
Forced expression of KLF4 cannot rescue fusion, *Myog* or *Npnt* expression in *Smad2*-deficient myoblasts. **(A)** Immunostaining for MyHC reveals myotubes generated from primary myoblasts isolated from *Smad2*^fl/fl^ mice retrovirally transduced to express CreER were treated with 4-OH tamoxifen to excise *Smad2* (TAM) or vehicle-treated (Veh), transduced to express KLF4 or with empty virus (EV) and differentiated for 2 days. Scale bar = 50 µm. **(B)** Differentiation index (# nuclei in MyHC+ cells/ total nuclei) from cells differentiated as in (A). n=4. **(C)** Fusion index (#nuclei/myotube) of cells cultured, transduced and differentiated as in (A). n=4. **(D)** RT-qPCR analysis of *Smad2, Klf4, Myog* and *Npnt* in cells transduced as in (A) and induced to differentiate for 6 h in low-serum conditions. n=4. **(E)** Transcription reporter assay measuring activity of the *Myog* promoter in C2C12 cells in the presence of SMAD2 and KLF4 relative to controls. n=5. **(F)** Reporter assay measuring activity from the *Myog* promoter in C2C12 cells transduced to express a shRNA directed against *Smad2* (shSmad2) in the presence or absence of KLF4. n=5. **(G)** Occupancy of the *Myog* promoter by SMAD2 and KLF4 by chromatin immunoprecipitation as compared to non-specific antibody (IgG). n=6 for SMAD2, n=3 for KLF4, ***p<0.001. **(H)** Coimmunoprecipitation of KLF4 with SMAD2 from whole cell extracts from differentiating C2C12 myoblasts 3 days after induction. IN is 10% of input used for IP. **(I)** Schematic representation of the regulatory region of the *Npnt* gene including putative SMAD2 and KLF4 motifs. **(J)** Chromatin immunoprecipitation of SMAD2 and KLF4 occupancy of 2 promoter regions in the *Npnt* promoter and -12 kb upstream region performed in C2C12 cells differentiated for 1 day in low serum conditions. Data are shown as copy numbers in comparison to pulldown with type-matched IgG as a control. n=3. Bars are means + STDEV.

To determine if SMAD2 and KLF4 cooperate to regulate *Npnt* expression, the *Npnt* gene regulatory region was analysed for putative KLF4 and SMAD binding sites. Primers were designed to amplify 3 specific regulatory regions of the *Npnt* gene: “pro1” which contains 1 KLF4 motif, “pro2” which has 2 KLF4 motifs and -12 kb region which contains 3 KLF4 motifs and 1 SMAD motif (Fig. 7I). Using publicly available ChIP-seq data, the -12 kb region was found to have H3K27Ac histone marks in myoblasts (GSE37525), corresponding with an active enhancer (Blum et al., 2012). Chromatin immunoprecipitation revealed that SMAD2 and KLF4 occupy all 3 *Npnt* regulatory regions examined under differentiation conditions in C2C12 myoblasts (Fig. 7J). Taken together these results suggest that SMAD2 is required for the regulation of *Npnt* by KLF4 during myogenic fusion.

### SMAD2 negatively regulates the expression of inhibitors of myogenic differentiation

To explore the molecular mechanism by which SMAD2 promotes myogenic differentiation, we performed an RT-qPCR array comparing the expression of 84 genes involved in myogenesis and myopathy in differentiating C2C12 myoblasts overexpressing SMAD2 and in primary myoblasts lacking SMAD2 along with their respective controls. We found 23 upregulated genes (≥1.5 fold change) in C2C12 cells overexpressing SMAD2 after 2 days of differentiation as compared to empty virus controls (Fig. 8A, cluster 1). As expected, the majority of upregulated genes were positive regulators of myogenic differentiation and maturation such as the structural genes *Acta1, Neb* (Nebulin) and the troponin genes (*Tnni2, Tnnc1, Tnnt1* and *Tnnt3*). In addition to these regulators, 18 genes were down regulated by ≥1.5 fold in cells overexpressing SMAD2 as compared to controls (Fig. 8A, cluster 2). Among these downregulated genes, we found known inhibitors of myogenic differentiation such as myostatin (*Mstn*), basic fibroblast growth factor (*Fgf2*), Bone morphogenetic protein 4 (*Bmp4*), interleukin 6 (*Il6*) and Mitogen-activated protein kinase 3 (*Mapk3*) (Fig. 8A, cluster 2). By comparison, knockout of SMAD2 in primary myoblasts resulted in the downregulation of 4 genes (*Myh2, Hk2, Myf6* and *Myf5*) at a threshold of ≥1.5 fold reduction as compared to control cells (Fig. 8A, cluster 3). Myosin heavy chain 2 (*Myh2*) is a marker of terminal differentiation, which is in line with the requirement of SMAD2 for efficient myogenic differentiation. Interestingly, 18 mRNAs were upregulated in *Smad2*^SC-/-^-derived primary myoblasts by at least 1.5-fold (Fig. 8A, cluster 4). Genes that failed to meet the significance cutoff are indicated in (B). Interestingly, only 5 genes were found among the genes downregulated in SMAD2-overexpressing cells and upregulated with loss of SMAD2 (Fig. 8A, genes in red). These genes, *Bmp4, Fgf2, Mstn, Igf1* and insulin growth factor binding protein 3 (*Igfbp3*) are, with the exception of *Igf1*, known potent inhibitors of myogenic differentiation. Taken together, these data suggest that SMAD2 regulates myogenic differentiation by inhibiting the expression of anti-myogenic factors that modulate the expression of myogenin and thus promotes myogenic differentiation.

**Figure 8.**
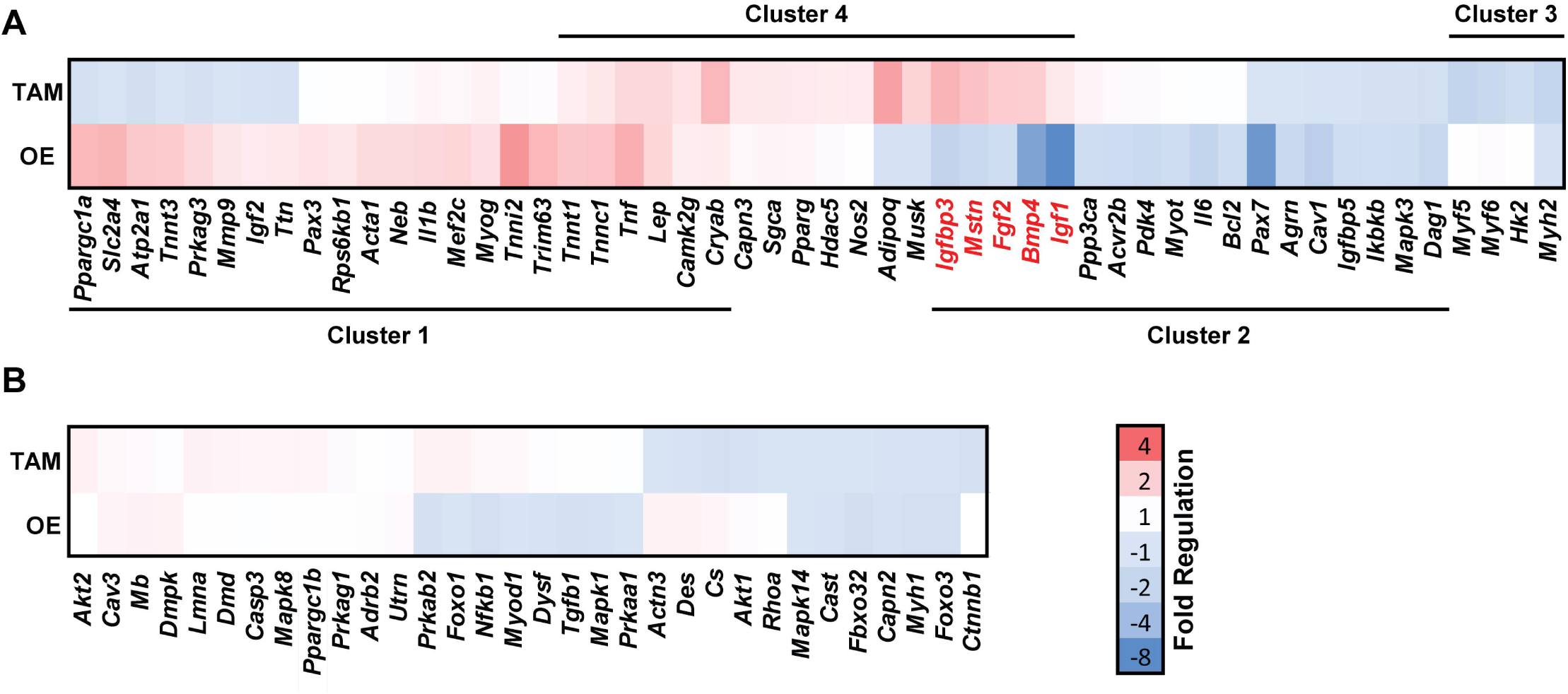
SMAD2 regulates inhibitors of myogenesis. C2C12 myoblasts retrovirally transduced to express SMAD2 (OE) and empty vector controls as well as primary myoblasts from *Smad2*^fl/fl^ mice retrovirally transduced to express CreER and treated with TAM or vehicle were induced to differentiate and isolated mRNA (pooled from 3 trials) was analyzed by RT2 Profiler Array (Qiagen) for 84 myogenesis and myopathy related genes. Heatmaps are shown as relative to controls for both OE and TAM displayed as fold-change. **(A)** Heatmap of differentially regulated genes by SMAD2 overexpression (OE) and SMAD2 knockout (TAM). Cluster 1 genes contains genes that are upregulated by ≥1.5-fold in C2C12 cells overexpressing SMAD2. Cluster 2 regroups genes that are downregulated by ≥1.5-fold with overexpression of SMAD2. Cluster 4 contains genes that are upregulated by ≥1.5-fold in Smad2^SC-/-^ (TAM). Cluster 4 are genes downregulated by ≥1.5-fold in Smad2^SC-/-^. Genes that are in red are upregulated with OE and downregulated in Smad2^SC-/-^ with a threshold of ≥1.5-fold. **(B)** Genes not significantly regulated in either condition.

## DISCUSSION

Herein, we identify SMAD2 as a powerful regulator of terminal myogenic differentiation and fusion. SMAD2 gain and loss of function experiments revealed that *Klf4, Myog* and *Npnt* are SMAD2 target genes. Our mRNA expression screen revealed that SMAD2 negatively regulates the expression of 5 genes: *Igf1, Igfbp3, Fgf2, Mstn* and *Bmp4*. While the mechanism by which SMAD2 inhibits the expression of these genes in skeletal muscle remains unknown, *Fgf2* was identified as a TGFβ target gene in stromal cells (Strand et al., 2014) and *Igfbp3* is a known TGFβ target during epithelial to mesenchymal transition (Natsuizaka et al., 2010). Interestingly, both IGF-1 and IGFBP3 can enhance activation of TGFβ receptors and can stimulate TGFβ activity (Fanayan et al., 2002; Kuemmerle et al., 2004; Natsuizaka et al., 2010; Peters et al., 2006; Rosendahl and Forsberg, 2006), suggesting that SMAD2 may act to inhibit classical TGFβ mediated responses. Indeed, coupled with the inhibition of both myostatin and BMP4, SMAD2 actions during myogenic differentiation effectively reduce both receptor activation as well as autocrine production of TGFβ family ligands. These findings raise the intriguing possibility that SMAD2 may function in two distinct pathways. In the absence of TGFβ or myostatin, SMAD2 acts in a pro-myogenic pathway, promoting regulators of late myogenesis while inhibiting the expression of potent inhibitors of myogenic differentiation (BMP4, *Mstn*) and stimulators of TGFβ action (IGF-1, IGFBP3). However, in the presence of TGFβ or myostatin, SMAD2 acts to inhibit myogenic differentiation. It remains unclear if SMAD2 is acting on different gene targets in these two proposed mechanisms, or rather its activity is altered by phosphorylation and complexing with SMAD4, resulting in unique gene expression patterns regulated by SMAD2 in the presence and absence of TGFβ.

The interaction with, and apparent cooperation with KLF4 may represent a mechanism by which SMAD2 activity can be modulated. Both SMAD2 and SMAD3 are weak DNA binders on their own, and given that SMAD binding motifs are very frequent, specificity and transcriptional activity appears to be mediated by interactions with other transcriptions factors such as SMAD4, and in skeletal muscle, MYOD (Mullen et al., 2011). As such, phosphorylation by the TGFβ receptor complex and interaction with SMAD4 is considered essential for occupancy of regulatory elements. Interestingly, mutations in or deletion of *Smad4* are found in many cancers, and in the absence of SMAD4 protein, a subset of TGFβ targets genes are unaffected, suggesting that SMAD4-independent gene transcription downstream of TGFβ occurs (Levy and Hill, 2005). In our system, effects of SMAD2 on myogenic differentiation are independent of the C-terminal serine residues targeted by the TGFβ type I receptor. Thus, during normal myogenic differentiation, SMAD2 recruitment to target genes involved in terminal differentiation and fusion may occur through interaction with lineage-specific factors such as MYOD and/or KLF4 much in the way that SMAD4 brings SMAD2 to gene targets. As such, activation of TGFβ receptors and SMAD2 C-terminal phosphorylation may divert SMAD2 activity towards anti-myogenic targets, and act as a switch between an anti-myogenic program and a pro-differentiation program. Consistent with this notion, TGFβ1 and TGFβ2 expression is highest in proliferating myoblasts where phosphorylated SMAD2 is detected, and decreases with differentiation, precisely when the pro-myogenic role described herein is active. This raises the possibility that SMAD2 phosphorylation marks proliferative cells that cannot differentiate, and that suppression of this phosphorylation allows SMAD2 to assume a pro-myogenic role, driving the expression of KLF4, *Myog* and *Npnt*.

To influence gene expression, SMAD2 must gain entry into the nucleus, a process that in the context of TGFβ signaling requires both phosphorylation of SMAD2 and its interaction with the co-SMAD, SMAD4. While TGFβ has been shown to regulate the interaction of SMAD2 with SMAD4 in a phosphorylation-dependent mechanism, the transcriptional output from SMAD2-dependent genes appears to be mediated more by the retention of phosphorylated SMAD2 in the nucleus, rather than its import (Schmierer and Hill, 2005). Indeed, TGFβ signaling does not appear to regulate the nuclear import rate for SMAD2, but rather decreases its export from the nucleus (Xu et al., 2002). However, phosphorylation of C-terminal serine residues by the ligand-bound TGFβ receptor is believed to induce a conformational change that allows both interaction with SMAD4 and more efficient interaction with DNA response elements in target promoters, a situation that is unlikely to happen in our current model. Given the known collaboration of SMAD3 with master transcription factors such as MYOD and OCT4, it remains possible that transcription factors such as MYOD could also recruit SMAD2 to promote regulation of their own target genes (Mullen et al., 2011).

## MATERIALS AND METHODS

### Mice and Animal Care

All animal work was performed in accordance with the guidelines set out by the Canadian Council on Animal Care and was approved by the University of Ottawa Animal Care Committee. Smad2^tm1.1Epb^ (Smad2^fl/fl^ mice) (Ju et al., 2006) were obtained from the Jackson Laboratory and crossed with mice bearing the Pax7-CreER™ (Pax7^CreER^) (Nishijo et al., 2009). Smad2^fl/fl^*Pax7*^*+/+*^ (control) and conditional null Smad2^-/-^Pax7^CreER-/+^ (Smad2^SC-/-^) mice were generated and activation of CreER™ *in utero* was achieved by a single gavage of 2.5 mg of tamoxifen (dissolved in corn oil) of pregnant dams when pregnancy was at E15.5 or by five daily i.p. injections of 1.5 mg of tamoxifen dissolved in corn oil. For injury experiments, one week after last dose of tamoxifen, mice were injured by injecting 30µl of 10µM cardiotoxin (Latoxan) to the left tibialis anterior (TA) muscle and TA muscles were harvested and sectioned 7 or 14 days post injury. All animals were housed in a controlled facility (22°C with 30% relative humidity on 12 hours light/dark cycle) and provided with food and water ad libitum.

### Constructs and reagents

MSCV CreERT2 puro was a gift from Tyler Jacks (Addgene plasmid # 22776) (Kumar et al., 2009). The LPCX-SMAD2 (Addgene plasmid #12636) and LPCX Smad2 deltaSSMS (Addgene plasmid # 12637) were gifts from Rik Derynck (Choy et al., 2000). pMXs-KLF4 was a gift from Dr. Toshio Kitamura (Addgene plasmid #13370) (Takahashi and Yamanaka, 2006) and the control vector pMXs-RFP, a gift from Dr. William Stanford. The pGL3-Klf4-Luc promoter was a kind gift from Dr. Christman at the Ohio State University (Karpurapu et al., 2014)(Karpurapu et al., 2014). The -2 kb myogenin-luc reporter construct was a gift from Dr. Alexandre Blais (Liu et al., 2010).

### Isolation of Primary Myoblasts and Cell Culture

Primary myoblasts from C57BL/6 and *Smad2* conditional knockout mice were obtained as previously described (Marchildon et al., 2012). Briefly, lower hind limb muscles from C57BL/6 mice aged 6-8 weeks were dissected and digested with collagenase (Roche). Isolated cells were plated on matrigel-coated dishes and allowed to grow in DMEM containing 20% fetal bovine serum (FBS) and 10% horse serum (HS) in the presence of 10ng/ml basic FGF and 2ng/ml HGF (Peprotech). Differentiation was induced when cells reached 70-80% confluence by culturing in DMEM containing 2% FBS and 2% HS (Differentiation medium, DM).

C2C12 cells were cultured in DMEM containing 10% FBS. To induce differentiation, growth medium was replaced with DMEM containing 2% HS, when cells were 80-90% confluent.

### Retroviral infection

Retrovirus for expression of SMAD2 or KLF4 was generated by retroviral expression plasmids or empty vector controls into Phoenix cells and virus was captured from supernatants after 2 days. For viral infection, growth medium was replaced with a medium containing virus when C2C12 cells or primary myoblasts reached 30%-40% confluency. 48 hours after infection, 2µM puromycin was added to culture medium to select positive cells.

### Western analysis

Western blotting was performed as described (Fu et al., 2015). Briefly, whole cell lysate of primary myoblasts was prepared by lysing buffer containing protease inhibitor cocktail (Roche). Protein concentration was determined with the BCA Protein Assay Reagent (Pierce Biotechnology) using Bovine Serum Albumin as standard. Samples were subjected to SDS-PAGE gels and transferred to PVDF membranes (Millipore). Proteins were detected by the following antibodies: anti-Smad2 antibodies (Cell signaling, 5339S), anti-GKLF (KLF4) antibodies (Santa Cruz BioTech, sc-20691), anti-Myogenin antibodies (DSHB, F5D), anti-cyclophilin B (Abcam, ab16045) and anti-αTubulin antibodies (Santa Cruz BioTech, sc-5286).

### Immunofluorescence

Immunofluorescence staining was performed as described (Marchildon et al., 2012). Cultured cells were fixed by ice-cold methanol and permeabilization with PBS containing 0.5% Triton X-100. Detection was performed following standard procedures by using anti-MF20 (DSHB) and anti-Mouse-Cy3 (Jackson ImmunoResearch) antibodies. The differentiation index (DI) of myoblasts and fusion index (FI) of myotubes were calculated as described (Lamarche et al., 2015). Cryosections were fixed with 4%PFA and processed as described before (Lala-Tabbert et al., 2016). Primary antibodies used were PAX7 (DSHB), MYOG (DSHB, F5D) and dystrophin (Abcam, ab15277) followed by the secondary anti-mouse-Cy3 and Alexa-488 (Jackson ImmunoResearch).

### Reporter Assay

C2C12 cells were transiently transfected with reporter construct, mammalian expression constructs and control Renilla plasmid using FuGENE HD transfection reagent according to manufacturer’s instructions. Cells were supplemented with growth medium for 6 hours and collected 48 hours post-transfection. Extracts were analyzed using the Dual-Luciferase Reporter Assay Kit (Promega). The ratio of Luciferase/Renilla was calculated and normalized to experimental control (promoter in absence of experimental plasmid).

### Quantitative PCR analysis

Cells were harvested and RNA was extracted using the RNA Easy Mini Kit (Qiagen Hilden, Germany) according to the manufacturer’s instructions. Quantification was performed using a Nanodrop (Thermo) and 1 ug RNA was treated with RNase-Free DNase. First strand cDNA was made using the iScript cDNA Synthesis Kit (Bio-Rad). CT values from quantitative PCR were analyzed with the delta delta CT method using 18S as an internal control (Livak and Schmittgen, 2001). The primer sequences used were as follows: *Smad2* (F: 5□-ATGTCGTCCATCTTGCCA-3□; R: 5□-AACCGTCCTGTTTTCTTTAG-3□), *Smad3* (F: 5□-CGTAATTCATGGTGGCTGTG-3□; R: 5□-ACCAAGTGCATTACCATCCC-3□), *Cebpb* (F: 5□-TCGAACCCGCGGACTGCAAG-3′; R: 5□-CGACGACGACGTGGACAGGC-3□), *Pax7* (F: 5□-GACGACGAGGAAGGAGACAA-3□; R: 5□-CGGGTTCTGATTCCACATCT-3□); *18s* (F: 5□-CGCCGCTAGAGGTGAAATC-3□; *18s* R: 5□-CCAGTCGGCATCGTTTATGG-3□), *Myod1* (F: 5□-TGGCATGATGGATTACAGCG-3□; R: 5□-CCACTATGCTGGACAGGCAGT-3□); *Myog* (F: 5□-ATCGCGCTCCTCCTGGTTGA-3□, R: 5□-CTGGGGACCCCTGAGCATTG-3□), *Myh8* (F: 5□-TCGCTGGCTTTGAGATCTTT-3□, R: 5□-ACGAACATGTGGTGGTTGAA-3□), *Klf4* (F: 5□-GCAGTCACAAGTCCCCTCTC-3□; R: 5□-TAGTCACAAGTGTGGGTGGC-3□), *Npnt* (F: 5□-TGGAGGCAAACCCAGATCAC-3; R: 5□-GCAGCGACCTCTTTTCAAGC-3□), *Tmem8c* (F: 5□-ATCGCTACCAAGAGGCGTT-3□; R: 5□-CACAGCACAGACAAACCAGG-3□) (Millay et al., 2013)

### Chromatin Immunoprecipitation

Chromatin Immunoprecipitation (ChIP) analysis was performed as previously described (Lala-Tabbert et al., 2016). Briefly, cells were cross-linked with 1% formaldehyde for 30 min at room temperature then sonicated for 30 cycles (30s on/ 30s off) with a Diagenode bioruptor. Equal amounts of chromatin were incubated with antibodies against KLF4-abcam, SMAD2 (abcam), or rabbit IgG as a negative control. Immunoconjugates were captured using protein G magnetic Dynabeads (Invitrogen), and DNA fragments were then purified with QIAquick PCR kit (Qiagen). A 10% input sample of each condition was used to generate a standard curve and the copy numbers of each immunoprecipitate is presented relative to the standard curve. Primer sequences and genome coordinates for qPCR-ChIP are as follows: *Klf4* pro1 F (chr4:55532503-55532525, mm10): 5′-TATAACTTCTCGCTCGCTTGCTC-3′ *Klf4* pro1 R (chr4:55532632-55532653, mm10): 5′-TGCGCGGAGTTTGTTTATTTAG-3′ *Klf4* -10kb F (Chr4:55542941-55542961, mm10): 5′-CAGGAATGCCTGTGGGGATAG-3′ *Klf4* -10kb R (Chr4:55543026-55543045, mm10): 5′-TGACCGGCTGAAGCTTTGTC-3′ *Myog* promoter F (chr1:134289663-134289681, mm10): 5′-GAGGCCCGGGTAGGAGTAA-3’ *Myog* promoter R (chr1:134289763-134289782, mm10): 5′-GCCGTCGGCTGTAATTTGAT-3’ *Npnt* pro1 F (chr3:132949265-132949284, mm10): 5′-GCTTTTCCTCTGGTCCCCTC-3′ *Npnt* pro1 R (chr3:132949336-132949355, mm10) : 5′-GATGCCGCACCTGTTTTACC-3′ *Npnt* pro2 F (chr3:132949284-132949303, mm10): 5′-ATTGGCTTTCCTGTCCCTGG-3′ *Npnt* pro2 R (chr3:132949107-132949126, mm10): 5′-TTAGCCTCTGGCTGCTTTCC-3′ *Npnt* -12kb F (chr3:132962692-132962711, mm10) : 5′-GGTTGGAACCGCAGTGAGTA-3′ *Npnt* -12kb R (chr3:132962807-132962826, mm10) : 5′-GCAGTGACAACAGGGAGACA-3′

### RT^2^ Profiler PCR Array

cDNA from Smad2 null and overexpressing cells and their relative controls was tested using the real-time RT^2^ Profiler PCR Array (QIAGEN, PAMM-099Z, Skeletal Muscle: Myogenesis and Myopathy RT^2^ Profiler PCR Array) in combination with SsoAdvanced Universal SYBR Green Supermix (Bio-Rad). Ct values were exported to an Excel file to generate a Ct value table. The table was then uploaded on to the data analysis web portal at http://www.qiagen.com/geneglobe and fold change/regulation was calculated using delta delta Ct method. Average of reference genes was used as internal control. Heatmap of fold regulation was prepared using Microsoft Excel.

### Statistical analysis

Statistical analysis was conducted using GraphPad Prism 6. A Student’s t-test was used when comparing two conditions. One-Way ANOVA was performed when comparing three or more treatments in one cell type. Two-way ANOVA was used when comparing two conditions in an experimental and control cell line. *Post-hoc* tests followed only statistically significant ANOVA results (p <0.001).

## ACKNOWLEDGEMENTS

The authors wish to thank James Haskins, François Marchildon, Dr. Qiao Li, Dr. David Lohnes, and Dr. Alexandre Blais for their input. We benefited from the use of several antibodies, listed in the “Methods” section, that were obtained from the Developmental Studies Hybridoma Bank (DSHB) developed under the auspices of the NICHD and maintained by The University of Iowa, Department of Biology, Iowa City, IA, USA. All animal work was performed following Canadian Council on Animal Care guidelines and was approved by the University of Ottawa Animal Care Committee.

